# Differential associations of insulin resistance with anterior hippocampal volume in mild cognitive impairment and Alzheimer’s disease

**DOI:** 10.64898/2026.08.04.742662

**Authors:** Nayu Watanabe, Akitoshi Ogawa, Takahiro Osada, Yusuke Adachi, Tomohiko Shirokoshi, Hiroyasu Kodama, Yasushi Oshima, Sakae Tanaka, Hideyoshi Kaga, Yoshifumi Tamura, Hirotaka Watada, Ryuzo Kawamori, Seiki Konishi

## Abstract

Insulin resistance is increasingly recognized as a metabolic factor associated with Alzheimer’s disease (AD); however, its relevance to hippocampal structural changes—a key pathological feature of AD—across disease stages is not fully understood. To address this issue, we investigated the relationship between insulin resistance, hippocampal gray matter volume, and cognitive performance using data from the Alzheimer’s Disease Neuroimaging Initiative (ADNI), a large-scale neuroimaging dataset. Insulin resistance was assessed using the Homeostatic Model Assessment of Insulin Resistance (HOMA-IR), and its relationship with brain structure and cognitive performance was evaluated across diagnostic groups. In the mild cognitive impairment (MCI) group, higher insulin resistance was associated with larger anterior hippocampal gray matter volume, whereas in the AD group this association was reversed in direction. Furthermore, in the MCI group, anterior hippocampal gray matter volume was also positively associated with higher Mini-Mental State Examination (MMSE) scores, and an exploratory mediation analysis suggested a significant indirect association linking HOMA-IR, anterior hippocampal volume, and cognitive performance through anterior hippocampal volume. These findings suggest that the relationship between insulin resistance and AD-related brain changes differs across diagnostic groups, highlighting the importance of considering metabolic alterations in relation to disease status.

## Introduction

Alzheimer’s disease (AD) is a progressive neurodegenerative disorder and the leading cause of dementia worldwide [1]. It is characterized by amyloid-β and tau pathologies that may begin years before overt dementia [2–5]. Clinically, AD-related changes are commonly described as progressing from preclinical phases through mild cognitive impairment (MCI) to AD dementia [6–9]. Hippocampal and medial temporal lobe atrophy are important structural markers of AD, as these changes are already evident at the MCI phase and are associated with cognitive decline and progression to AD dementia [10–13]. The hippocampus is central to episodic memory and is heterogeneous along its longitudinal axis, with the anterior hippocampus showing distinct functional specialization and AD-related structural vulnerability [14–16].

Metabolic dysfunction, particularly insulin resistance, has been increasingly implicated in cognitive decline and AD. Epidemiological studies have linked diabetes to an increased risk of dementia, and disrupted insulin signaling may contribute to AD-related pathology through impaired amyloid-β clearance, tau hyperphosphorylation, oxidative stress, and altered synaptic function [17–21]. However, human neuroimaging studies linking insulin resistance to AD-related brain alterations have yielded inconsistent findings, including altered medial temporal metabolism in individuals with MCI and reduced gray matter volume in AD-sensitive, default-mode, or limbic regions [22–24].

One possible explanation for these inconsistencies is that the association between insulin resistance and brain structure differs by disease status. The relationship between insulin resistance and hippocampal structure may therefore not be uniform across AD-related diagnostic groups. This possibility may be particularly relevant to the anterior hippocampus, which shows functional specialization and AD-related structural vulnerability.

In this study, we used data from the Alzheimer’s Disease Neuroimaging Initiative (ADNI) to examine whether the association between insulin resistance and anterior hippocampal gray matter volume differs across diagnostic groups. Insulin resistance was assessed using the Homeostatic Model Assessment of Insulin Resistance (HOMA-IR), a widely used surrogate index derived from fasting insulin and glucose levels [25]. Gray matter volume was evaluated using voxel-based morphometry (VBM) applied to structural MRI data, which enables voxel-wise assessment of gray matter alterations [26,27]. We further examined whether anterior hippocampal structure was associated with global cognitive performance, as assessed by the Mini-Mental State Examination (MMSE).

## Results

The MRI analysis included 526 ADNI participants classified as cognitively normal (CN), MCI, or AD (Fig. 1). We first summarized clinical characteristics across diagnostic groups and then examined group-specific associations of log-transformed HOMA-IR with anterior hippocampal gray matter volume and MMSE scores.

**Figure 1.**
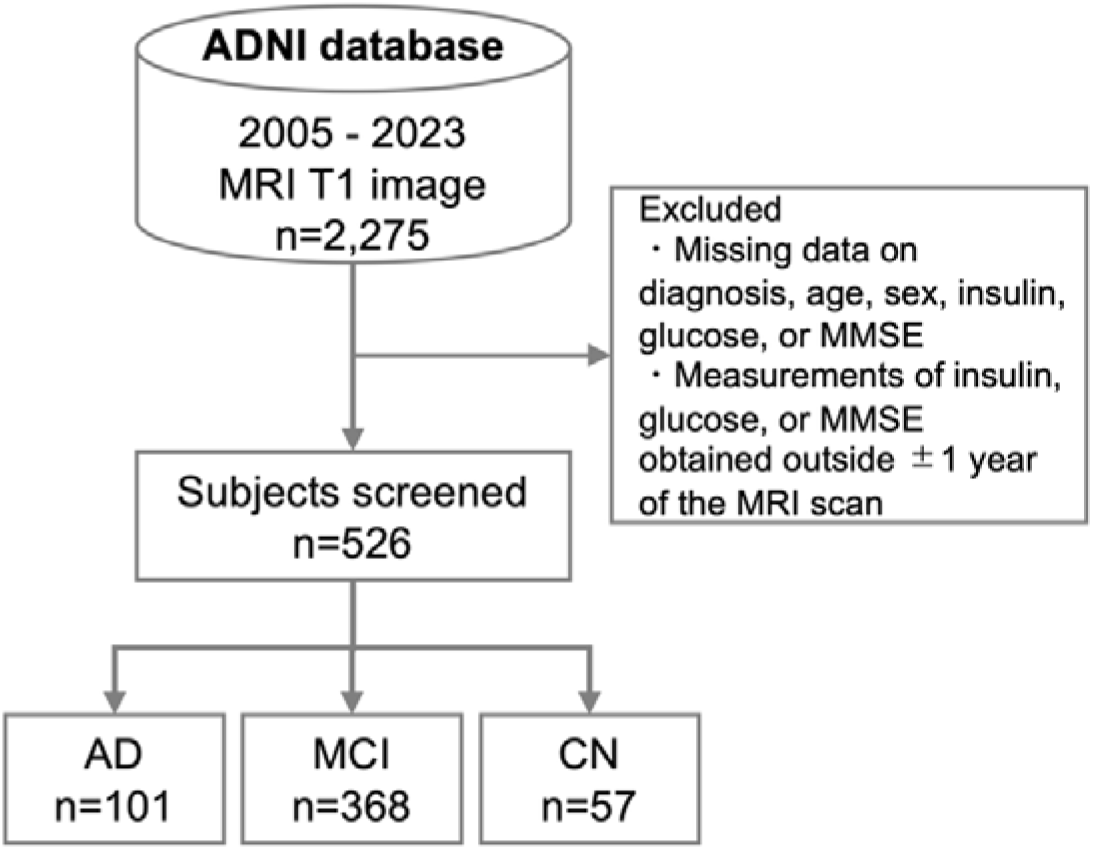
Flowchart of participant selection. Participants were selected from the ADNI database for the MRI analysis. Individuals were excluded if data on diagnostic group, age, sex, fasting insulin, fasting glucose, or Mini-Mental State Examination (MMSE) were missing, or if insulin, glucose, or MMSE measurements were obtained more than 1 year apart from the MRI scan. The final sample included 526 participants: 101 with Alzheimer’s disease (AD), 368 with mild cognitive impairment (MCI), and 57 cognitively normal (CN) individuals.

### Participant Characteristics

Participant characteristics are summarized in Table 1. Mean age was comparable across diagnostic groups. Fasting glucose, fasting insulin, and HOMA-IR values were broadly similar among groups. MMSE scores decreased from CN to MCI and AD, consistent with the diagnostic classification. Total brain volume (TBV) tended to be lower in the AD group than in the CN and MCI groups.

**Table 1.**
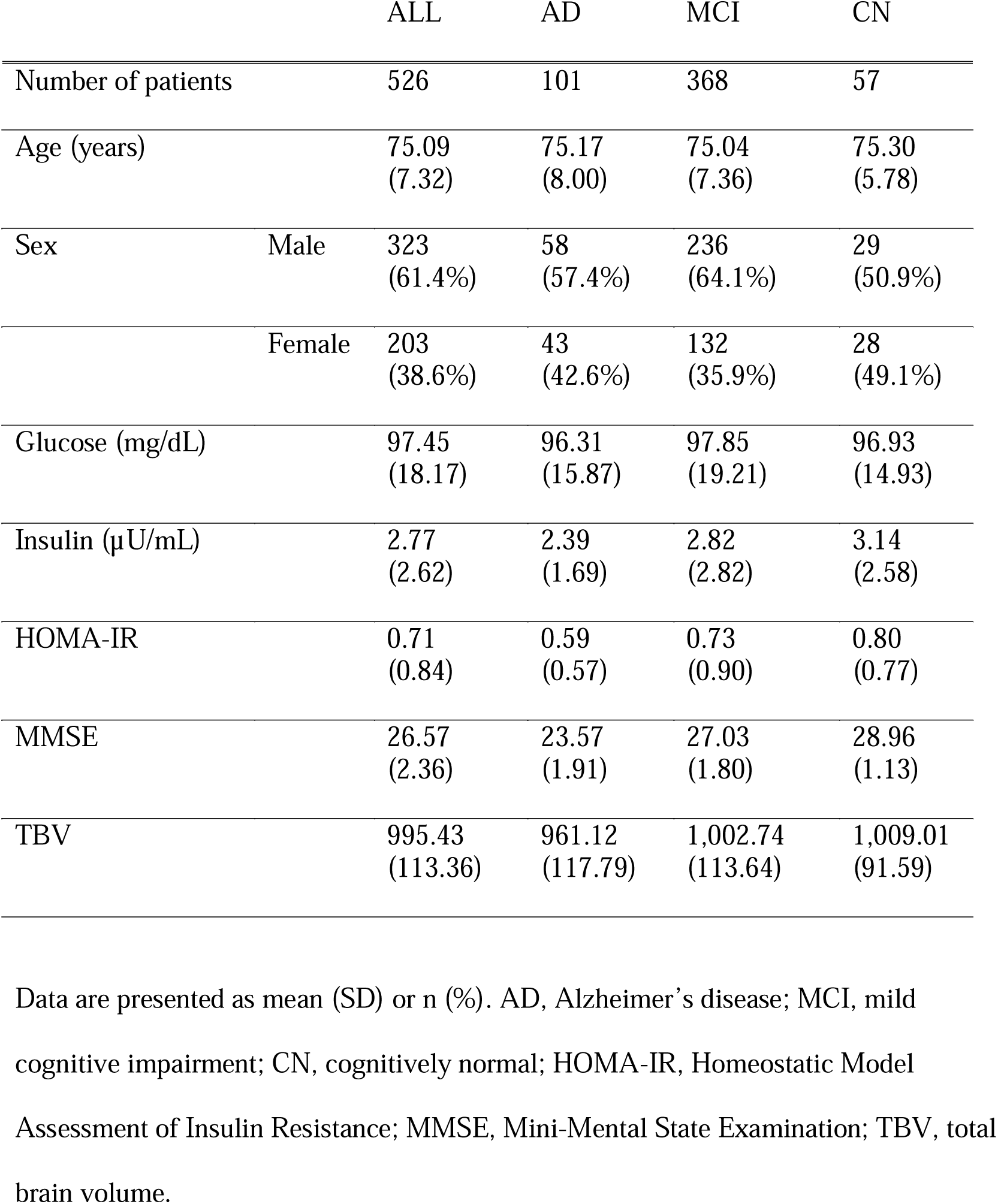
Clinical characteristics of the study participants.

Data are presented as mean (SD) or n (%). AD, Alzheimer’s disease; MCI, mild cognitive impairment; CN, cognitively normal; HOMA-IR, Homeostatic Model Assessment of Insulin Resistance; MMSE, Mini-Mental State Examination; TBV, total brain volume.

### VBM Results

Voxel-wise analyses were performed to identify brain regions in which gray matter volume was associated with HOMA-IR or MMSE within each diagnostic group. In the MCI group, HOMA-IR showed significant positive associations with gray matter volume in the medial temporal lobe, particularly within the bilateral anterior hippocampus (Fig. 2A). For this association, the peak coordinates of the major clusters were located at Montreal Neurological Institute (MNI) coordinates X = −33, Y = −15, Z = −16 in the left hemisphere and X = 34, Y = −15, Z = −16 in the right hemisphere, with peak-level family-wise error (FWE)-corrected p values using small volume correction (SVC) of 0.034 and 0.030, respectively. In the AD group, HOMA-IR exhibited a negative direction of association with gray matter volume in the medial temporal lobe, including the anterior hippocampus, in contrast to the positive association observed in the MCI group (Fig. 2B). Although this association did not reach statistical significance, the statistical map showed a spatial distribution within the anterior hippocampus that was opposite in direction to that observed in the MCI group.

**Figure 2.**
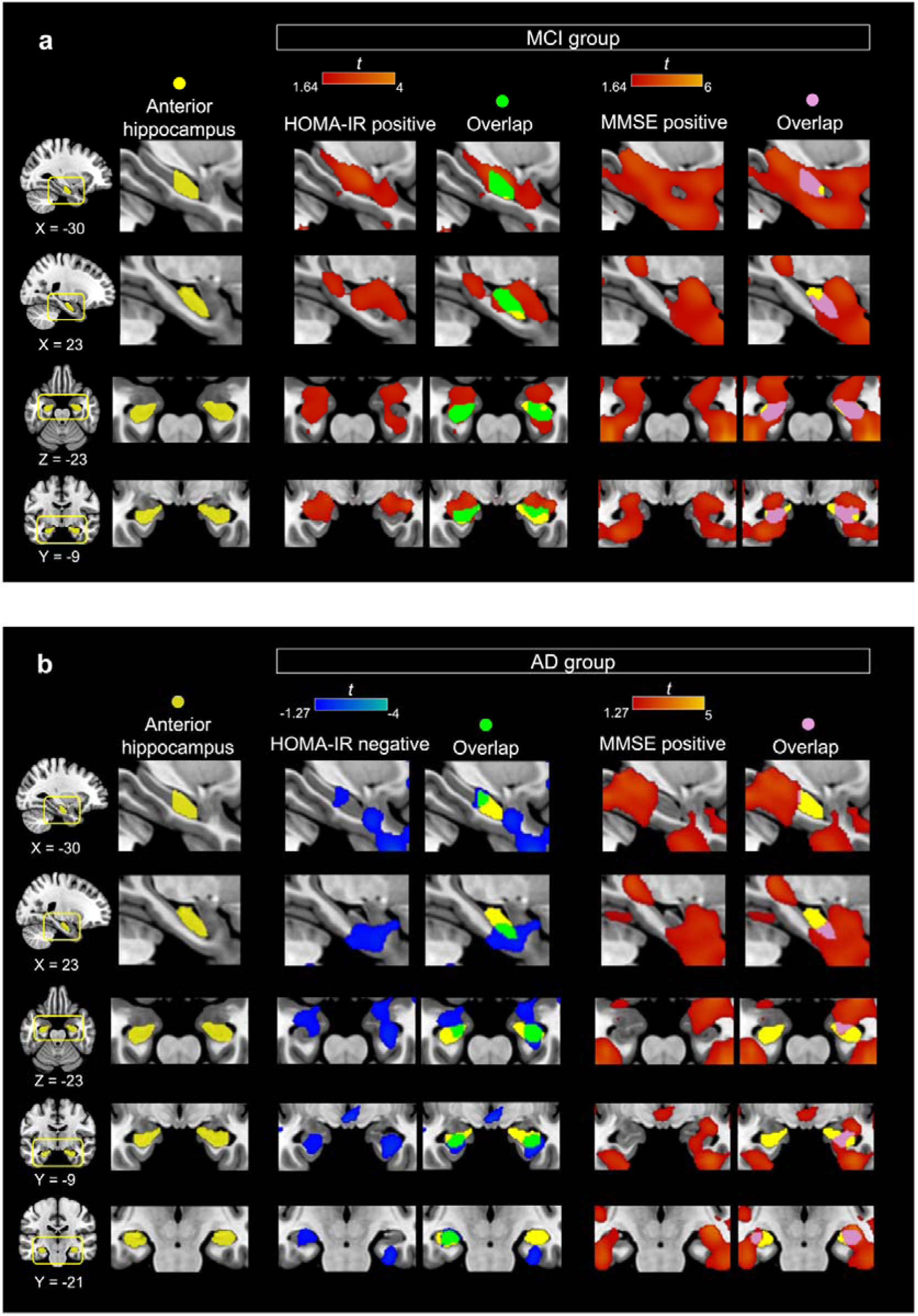
Voxel-based morphometry results for the anterior hippocampus. (a) In the MCI group, higher HOMA-IR was positively associated with gray matter volume (GMV) in the bilateral anterior hippocampus, and MMSE scores were positively associated with GMV in overlapping regions. (b) In the AD group, higher HOMA-IR was negatively associated with GMV in the anterior hippocampus, whereas MMSE scores were positively associated with GMV in partially overlapping regions. Warm colors indicate positive associations, whereas cool colors indicate negative associations. Yellow indicates the anterior hippocampal region of interest (ROI). Green and pink indicate overlap between the anterior hippocampal ROI and regions associated with HOMA-IR and MMSE, respectively. Images are displayed at an uncorrected threshold for visualization.

MMSE scores showed positive associations with gray matter volume in the anterior hippocampal regions in both diagnostic groups. In the MCI group, MMSE-associated clusters spatially overlapped with the HOMA-IR-associated clusters within the anterior hippocampus. In the AD group, MMSE scores remained positively associated with gray matter volume in these regions, whereas the HOMA-IR association was negative in direction. A visual overlay indicated partial spatial correspondence between the HOMA-IR pattern and MMSE-related regions within the anterior hippocampus.

No significant associations were observed in the CN group, although the smaller sample size in this group limits the statistical power to detect such associations. Whole-brain analyses are presented in Supplementary Figs. S1 and S2.

### Anterior Hippocampal ROI Analysis Results

To quantitatively evaluate anterior hippocampal involvement indicated by the VBM analyses, regression analyses were performed using gray matter volumes extracted from the anterior hippocampal region of interest (ROI) (Fig. 3). In group-stratified analyses, a positive association between HOMA-IR and anterior hippocampal gray matter volume was observed in the MCI group. In contrast, the AD group exhibited a negative direction of association, whereas no significant association was observed in the CN group. In the interaction model including all three diagnostic groups, the HOMA-IR × diagnostic group term comparing MCI with AD was significant (p = 0.017), indicating that the slope of the association between HOMA-IR and anterior hippocampal gray matter volume differed between the MCI and AD groups.

**Figure 3.**
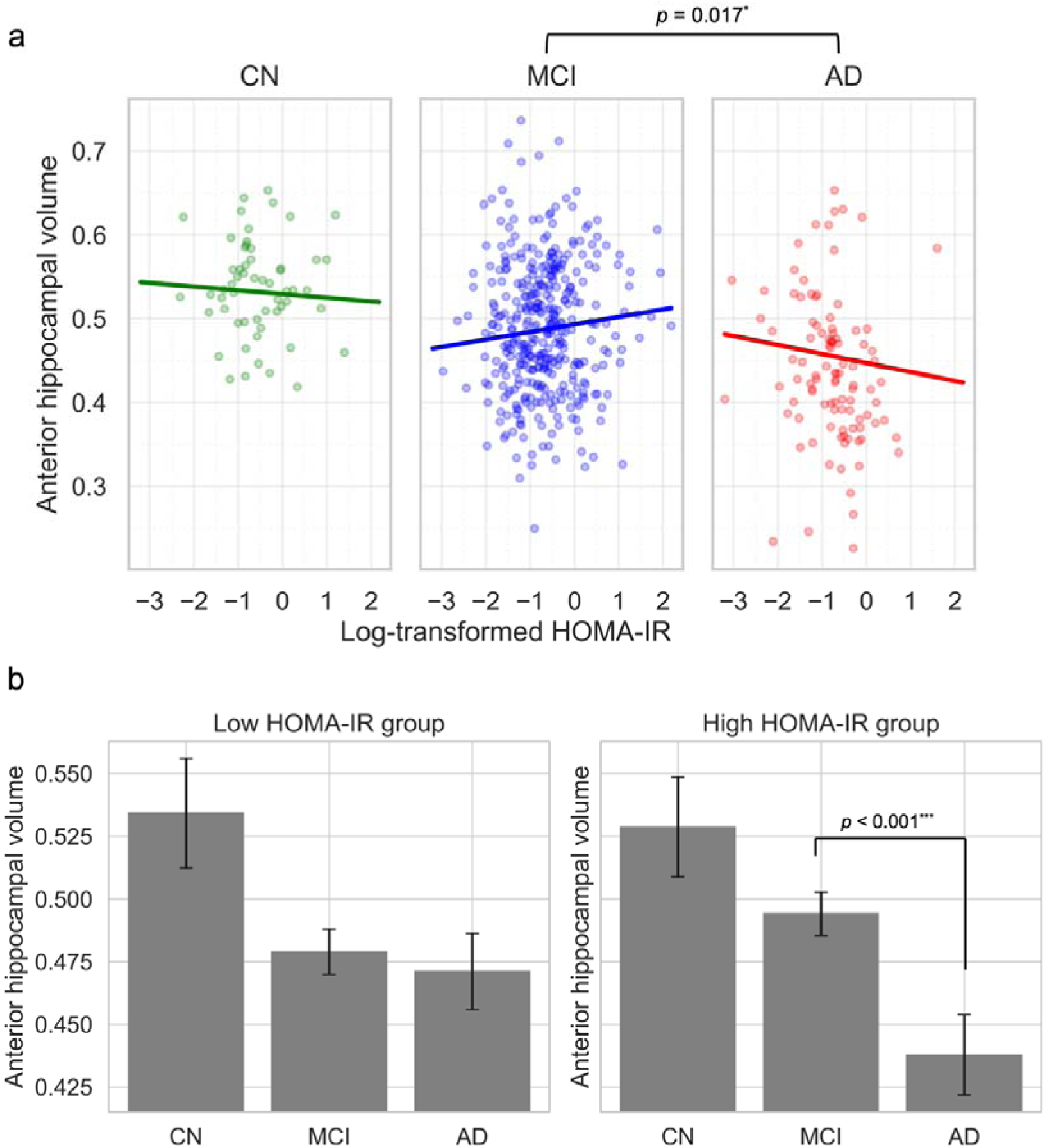
Group-specific associations between HOMA-IR and anterior hippocampal gray matter volume. (a) Associations between log-transformed HOMA-IR and anterior hippocampal gray matter volume (GMV) in each diagnostic group. A significant group × HOMA-IR interaction was observed between the MCI and AD groups (p = 0.017). (b) Anterior hippocampal GMV stratified by low and high HOMA-IR groups. In the high HOMA-IR subgroup, anterior hippocampal GMV was significantly lower in the AD group than in the MCI group (p < 0.001). No significant between-group difference was observed in the low HOMA-IR subgroup.

To further clarify the nature of the group × HOMA-IR interaction, additional analyses were conducted by stratifying participants into low- and high-HOMA-IR subgroups based on the median value of log-transformed HOMA-IR. In the high-HOMA-IR subgroup, anterior hippocampal gray matter volume was significantly lower in the AD group compared with the MCI group based on an age-, sex-, and TBV-adjusted linear model with Holm-corrected pairwise comparisons (adjusted p < 0.001). In contrast, no significant difference between the MCI and AD groups was observed in the low-HOMA-IR subgroup.

### Mediation Analysis Results

To further characterize the cognitive relevance of the anterior hippocampal association observed in the MCI group, we performed an exploratory mediation analysis examining whether anterior hippocampal gray matter volume statistically linked HOMA-IR and MMSE score (Fig. 4). In this model, HOMA-IR was significantly positively associated with anterior hippocampal gray matter volume (Path *a*), and anterior hippocampal gray matter volume was also significantly positively associated with MMSE score (Path *b*).

**Figure 4.**
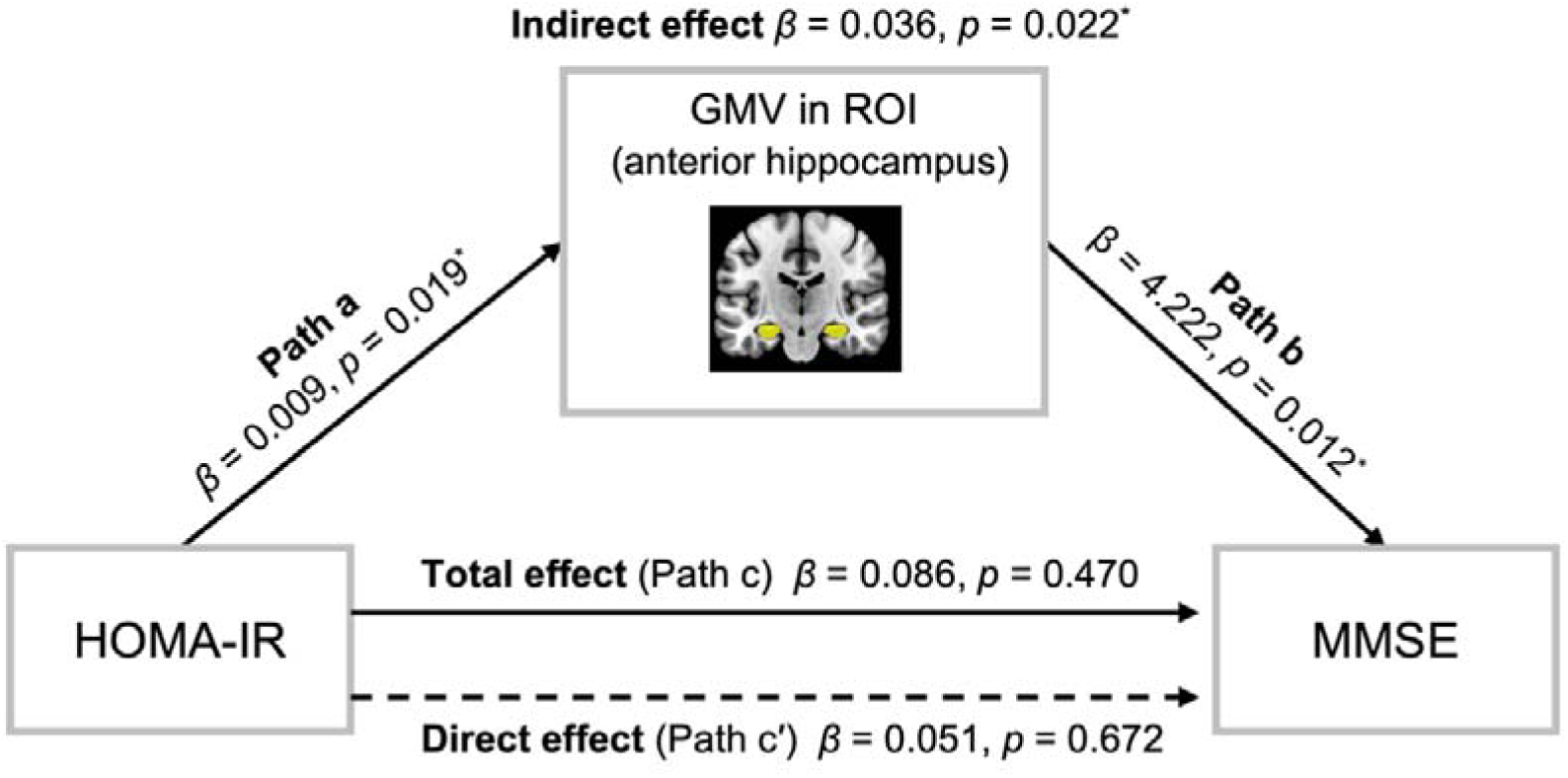
Mediation model of the association between HOMA-IR and MMSE through anterior hippocampal gray matter volume in the MCI group. Anterior hippocampal gray matter volume (GMV) was tested as a mediator of the association between log-transformed HOMA-IR and MMSE score. Path coefficients, total effect, direct effect, and indirect effect are shown in the figure. The indirect effect through anterior hippocampal GMV was significant, whereas the total and direct effects were not.

The indirect effect was statistically significant (p = 0.022), whereas neither the direct effect nor the total effect of HOMA-IR on MMSE reached statistical significance. These results suggest an indirect statistical association between HOMA-IR and cognitive performance through anterior hippocampal gray matter volume in the MCI group. Detailed mediation results for the anterior, posterior, and whole hippocampus are provided in Supplementary Table S1.

## Discussion

Our findings reveal that the association between insulin resistance (HOMA-IR) and anterior hippocampal gray matter volume is reversed across diagnostic groups. Specifically, higher HOMA-IR was associated with larger anterior hippocampal volume in the MCI group, whereas this relationship shifted to a negative direction in the AD group. This divergence was further supported by a significant interaction between HOMA-IR and diagnostic group. Notably, the brain regions in which HOMA-IR was associated with gray matter volume spatially overlapped with those associated with cognitive performance (MMSE scores) within the anterior hippocampus. Furthermore, exploratory mediation analysis in the MCI group suggested a statistically significant indirect pathway linking insulin resistance, anterior hippocampal volume, and cognitive function, although the direct and total effects were not statistically significant. These results suggest that the effect of insulin resistance on anterior hippocampal structure differs by disease status, indicating that the effect of metabolic dysfunction in Alzheimer’s disease is not uniform.

Several limitations should be considered when interpreting these findings. First, the cross-sectional design precludes direct examination of causal dynamics over time. Second, the sample size for participants with available insulin measurements was limited. Third, potential confounders, including diet and physical activity, were not fully accounted for. Fourth, future investigations incorporating hippocampal subfield-level analyses (e.g., CA1) may provide more granular insights. Most importantly, the present study did not directly evaluate the relationship between insulin resistance and molecular biomarkers using PET imaging, due to the limited number of participants with PET imaging data. As hippocampal atrophy is associated with both amyloid-β and tau pathologies, future research integrating PET imaging is essential to determine whether insulin resistance influences neurodegeneration independently of, or in interaction with, these pathologies.

The localization of the observed association to the anterior hippocampus further highlights the structural and functional heterogeneity of the hippocampus. The hippocampus is not a uniform structure and shows anatomical variation across subfields [28]. It also exhibits functional specialization along its longitudinal axis [15,29]. In particular, the anterior hippocampus is thought to differ functionally from posterior regions and to show distinct patterns of large-scale connectivity along the longitudinal axis [15,29]. Moreover, structural alterations in MCI are not evenly distributed across hippocampal subregions, and CA1 has been reported to be sensitive to structural changes at the predementia stage [30]. Our finding that the association with insulin resistance was localized to the anterior hippocampus, with a peak in a region corresponding to CA1, may therefore be understood in the context of this internal hippocampal heterogeneity.

In the anterior hippocampal region, the positive association between insulin resistance and gray matter volume observed in the MCI group may be interpreted in light of functional hyperactivity in the medial temporal lobe during the early phase of AD. Increased neural activity in the hippocampus has been reported in individuals with MCI compared with cognitively normal older adults [31]. Furthermore, Willette et al. [22] demonstrated that insulin resistance was associated with hypermetabolism in these brain regions in individuals with MCI. Our results further show that insulin resistance is associated with larger anterior hippocampal gray matter volume in MCI, extending prior findings from metabolism to structure. From a longitudinal perspective, increased hippocampal activity in Aβ-positive MCI has been reported to be associated with subsequent hippocampal atrophy and to predict clinical progression [32]. In this context, a transient state related to hyperactivity may underlie the positive association observed in the MCI group in our study.

Some previous studies have reported negative associations between insulin resistance or glycemic control and the hippocampal structure. Rasgon et al. [33] reported that higher insulin resistance was associated with reduced total hippocampal volume in cognitively normal middle-aged women at genetic risk for AD. Kerti et al. [34] showed that higher HbA1c levels were associated with lower memory performance, and that this relationship was partly mediated by hippocampal microstructure in healthy older adults. Notably, these associations have been observed even in cognitively normal individuals, suggesting that the effects of metabolic dysfunction may emerge relatively early. These findings contrast with our observations in the MCI group but are consistent with those in the AD group, indicating that the relationship between insulin resistance and the hippocampus may vary depending on the clinical context.

Brain insulin resistance may also be relevant when considering the relationship between peripheral insulin resistance and brain changes. Kullmann et al. [35] reviewed evidence that insulin action in the brain may influence both metabolism and cognition, positioning brain insulin resistance as a potential link between metabolic and cognitive disorders in humans. In addition, insulin transport across the blood-brain barrier has been suggested to be reduced in humans with insulin resistance [36]. Furthermore, in patients with AD, insulin and IGF-1 signaling are disrupted in the brain, particularly in the hippocampal formation. This disruption includes dysregulation of insulin receptor substrate-1 (IRS-1) and has been linked to cognitive decline [37]. However, the mechanisms linking peripheral metabolic dysfunction to insulin action in the brain, as well as the direction of causality, remain under debate.

In summary, the association between insulin resistance and anterior hippocampal structure varies across diagnostic groups, showing opposite directions of association, with a positive association in MCI and a negative direction of association in AD. These findings underscore the need to interpret this association in the context of disease status. Moreover, the positive association observed in MCI should not be interpreted as evidence of neuroprotection, but may reflect a transient phase preceding subsequent neurodegeneration in AD. Further studies incorporating longitudinal designs and molecular biomarker assessments are needed to clarify the temporal and pathological mechanisms underlying this relationship.

## Methods

### Study Design and Participants

We conducted a cross-sectional analysis using data from the ADNI. ADNI is a large multicenter longitudinal study in the United States that collects standardized clinical assessments, neuroimaging data, genetic information, and fluid biomarkers to investigate the progression of Alzheimer’s disease [38]. The ADNI study was approved by the institutional review boards of all participating institutions as detailed at https://adni.loni.usc.edu/wp-content/uploads/how_to_apply/ADNI_Acknowledgement_List.pdf. All procedures were conducted in accordance with the ADNI protocol. Written informed consent was obtained from all participants. Participants were classified as CN, MCI, or AD according to the ADNI diagnostic criteria.

The participant selection procedure is summarized in Fig. 1. We included T1-weighted MRI scans acquired between 2005 and 2023. Participants were excluded if data were missing for diagnostic status, age, sex, fasting insulin, fasting glucose, or Mini-Mental State Examination (MMSE) score. To ensure temporal consistency, eligible observations were defined as combinations in which MRI acquisition, blood sampling, and MMSE assessment occurred within a 365-day interval. Because fasting insulin measurements were available only for a subset of the ADNI cohort, blood sampling dates were used as the reference time point for linking MRI scans, MMSE assessments, and diagnostic information. For participants with multiple eligible observations, one observation per participant was selected. When multiple eligible observations were available for a participant, the observation was selected first according to diagnostic priority in the order of AD, MCI, and CN, and then according to the shortest interval among MRI acquisition, blood sampling, and MMSE assessment. Diagnostic status was assigned based on ADNI diagnostic records around the blood sampling date. No participants were excluded after MRI quality control. Accordingly, the final sample size was primarily determined by availability of fasting insulin measurements. Applying these criteria, 526 participants were included in the final MRI analyses (CN, n = 57; MCI, n = 368; AD, n = 101).

### Clinical Measures

Clinical assessments and blood test data collected as part of the ADNI protocol were used in this study. Demographic variables included age and sex. Cognitive function was assessed using the MMSE, a standardized 30-point screening test for global cognitive function [39]. The MMSE is a copyrighted instrument and may not be used or reproduced in whole or in part, in any form or language, or by any means without written permission of PAR (www.parinc.com). MMSE scores were used to characterize cognitive performance and to examine associations with brain structural measures, with higher scores indicating better cognitive function.

HOMA-IR, an index of insulin resistance, was calculated from fasting insulin (µU/mL) and fasting glucose (mg/dL) levels using the following formula:

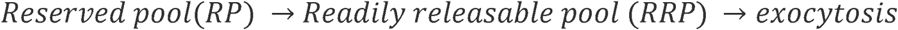

Because HOMA-IR values exhibited a right-skewed distribution, logarithmic transformation was applied prior to statistical analyses to improve conformity with the assumptions of linear modeling.

### MRI Data and Preprocessing

T1-weighted structural MRI data obtained from the ADNI database were used in this study. High-resolution three-dimensional T1-weighted images were obtained according to standardized ADNI acquisition procedures. All MRI preprocessing was performed using Statistical Parametric Mapping software (SPM12) [40]. Images were segmented into gray matter, white matter, and cerebrospinal fluid, followed by spatial normalization using the Diffeomorphic Anatomical Registration Through Exponentiated Lie Algebra (DARTEL) algorithm [41] and resampling to 1.5-mm isotropic voxels.

Normalized gray matter images were smoothed with an 8-mm full-width at half-maximum Gaussian kernel to reduce spatial noise. Modulated gray matter images, corrected for local volume changes introduced during nonlinear normalization, were used for subsequent structural analyses. TBV was derived from MRI preprocessing. It was included as a covariate in neuroimaging analyses to account for inter-individual differences in global brain size. Quality control procedures were applied throughout the MRI preprocessing pipeline, and no participants were excluded due to visible artifacts or processing errors.

### Neuroimaging Analysis

VBM analyses were performed using SPM12 to investigate associations between insulin resistance, cognitive function, and brain structure at the whole-brain voxel-wise level. Log-transformed HOMA-IR values were used in all analyses. A single general linear model (GLM) was constructed using modulated, normalized gray matter images pooled across all participants. The design matrix included diagnostic group (CN, MCI, and AD) and group-specific HOMA-IR regressors, allowing the association between HOMA-IR and gray matter volume to be evaluated within each diagnostic group. Age, sex, and TBV were included as covariates. Statistical significance was assessed using SVC within a bilateral hippocampal mask derived from the Neuromorphometrics atlas included in SPM12, applying FWE correction at the peak level (p < 0.05). To examine associations between cognitive function and brain structure, an additional single GLM was constructed using group-specific MMSE regressors, with the same covariate adjustment and statistical threshold.

ROI analyses focused on the anterior hippocampus. Hippocampal masks were generated from the Neuromorphometrics atlas included in SPM12. The hippocampus was subdivided into anterior and posterior regions along the longitudinal axis in MNI space, using a boundary at y = −21, based on prior studies [15]. Mean gray matter values were extracted separately for the left and right anterior hippocampus, and their average was used in subsequent analyses. Linear regression models were used to examine the association between insulin resistance and anterior hippocampal gray matter volume. Anterior hippocampal gray matter volume was used as the dependent variable, with log-transformed HOMA-IR as the independent variable. To determine whether the HOMA-IR–gray matter volume association differed across diagnostic groups, the models included diagnostic group (CN, MCI, and AD), the main effect of HOMA-IR, and the HOMA-IR × diagnostic group interaction term. Age, sex, and TBV were included as covariates. In addition, to further characterize the association, participants were stratified into high- and low-HOMA-IR groups based on the median value of HOMA-IR, and ROI analyses were performed within each group.

### Mediation Analysis

Mediation analyses were conducted to examine whether anterior hippocampal gray matter volume mediated the association between insulin resistance and cognitive function. HOMA-IR was specified as the independent variable, anterior hippocampal gray matter volume as the mediator, and MMSE score as the dependent variable. Age, sex, and TBV were included as covariates. The indirect effect was assessed using a bootstrap approach with 1,000 resampling iterations. ROI and mediation analyses were conducted using Python.

### Use of generative AI

During the course of preparing this work, the authors used Google Gemini for the purpose of language editing and manuscript refinement. Following the use of this service, the authors formally reviewed the content for its accuracy and edited it as necessary. The authors take full responsibility for all the content of this publication.

### Data Availability

The data analyzed in this study are available from the corresponding authors upon reasonable request to the extent permitted by ADNI data use policies. The original ADNI data are available from ADNI but are not publicly available from the authors.

## Acknowledgements

Data used in this study were obtained from the Alzheimer’s Disease Neuroimaging Initiative (ADNI) database (adni.loni.usc.edu). As such, the investigators within the ADNI contributed to the design and implementation of ADNI and/or provided data but did not participate in analysis or writing of this manuscript. A complete listing of ADNI investigators can be found at: http://adni.loni.usc.edu/wp-content/uploads/how_to_apply/ADNI_Acknowledgement_List.pdf

## Author Contributions

N.W. analyzed the data, interpreted the results, and drafted the manuscript. A.O., T.O., and S.K. contributed to the study design and provided input on the data analysis and interpretation. Y.A., T.S., H.Ko., Y.O., S.T., H.Ka., Y.T., H.W., and R.K. reviewed the manuscript and contributed to interpretation of the results. All authors approved the final version of the manuscript.

### Additional Information Competing interests

The authors declare no competing interests.

