## Supplementary for "Differential associations of insulin resistance with anterior hippocampal volume in mild cognitive impairment and Alzheimer’s disease"

**Supplementary Materials**

**Supplementary Table 1. Mediation analysis of the association between HOMA-IR and MMSE via hippocampal gray matter volume (GMV).**

| **Mediator (ROI)** | **Path / Effect** | **β** | **SE** | **95% CI (lower)** | **95% CI (upper)** | **p-value** |
| --- | --- | --- | --- | --- | --- | --- |
| **Anterior hippocampus** | a (HOMA-IR → GMV) | 0.009 | 0.004 | 0.001 | 0.016 | 0.019 |
|  | b (GMV → MMSE) | 4.223 | 1.674 | 0.932 | 7.514 | 0.012 |
|  | Direct effect (c′) | 0.051 | 0.119 | -0.184 | 0.285 | 0.672 |
|  | Indirect effect (a×b) | 0.036 | 0.021 | 0.006 | 0.091 | 0.022 |
| **Posterior hippocampus** | a (HOMA-IR → GMV) | 0.005 | 0.003 | 0 | 0.01 | 0.073 |
|  | b (GMV → MMSE) | 7.741 | 2.308 | 3.202 | 12.279 | <0.001 |
|  | Direct effect (c′) | 0.05 | 0.118 | -0.183 | 0.282 | 0.674 |
|  | Indirect effect (a×b) | 0.037 | 0.024 | 0.001 | 0.099 | 0.06 |
| **Whole hippocampus** | a (HOMA-IR → GMV) | 0.007 | 0.003 | 0.001 | 0.013 | 0.017 |
|  | b (GMV → MMSE) | 6.039 | 2.001 | 2.103 | 9.974 | 0.003 |
|  | Direct effect (c′) | 0.043 | 0.119 | -0.191 | 0.276 | 0.720 |
|  | Indirect effect (a×b) | 0.044 | 0.024 | 0.009 | 0.103 | 0.008 |

β, regression coefficient; SE, standard error; CI, confidence interval. Indirect effects were estimated as a×b.

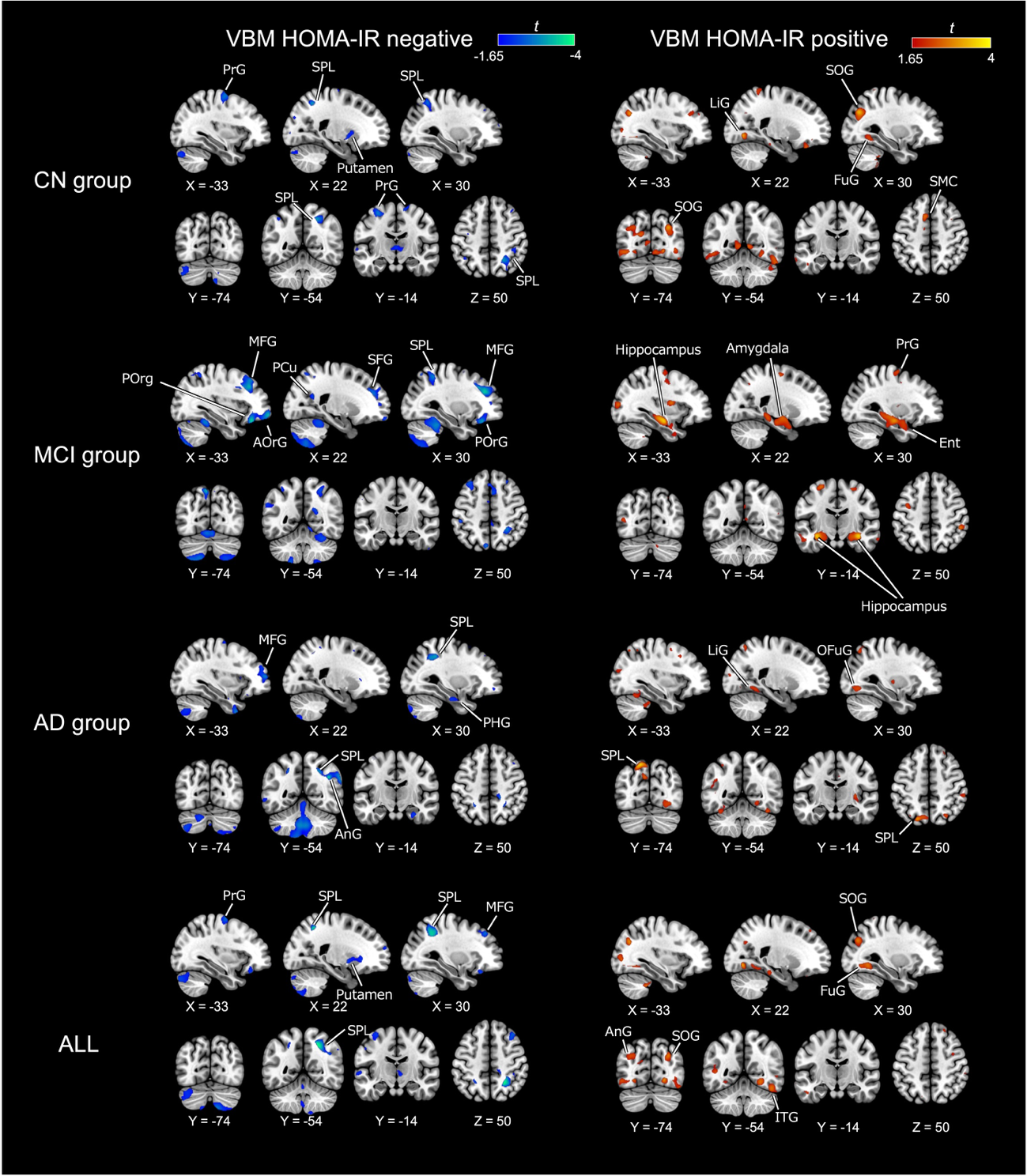
**Supplementary Figure 1. Voxel-wise associations between HOMA-IR and gray matter volume across diagnostic groups.**

Regions showing positive (warm colors) and negative (cool colors) associations with HOMA-IR are displayed for CN, MCI, AD, and all participants. Statistical maps are shown at an uncorrected threshold for visualization.

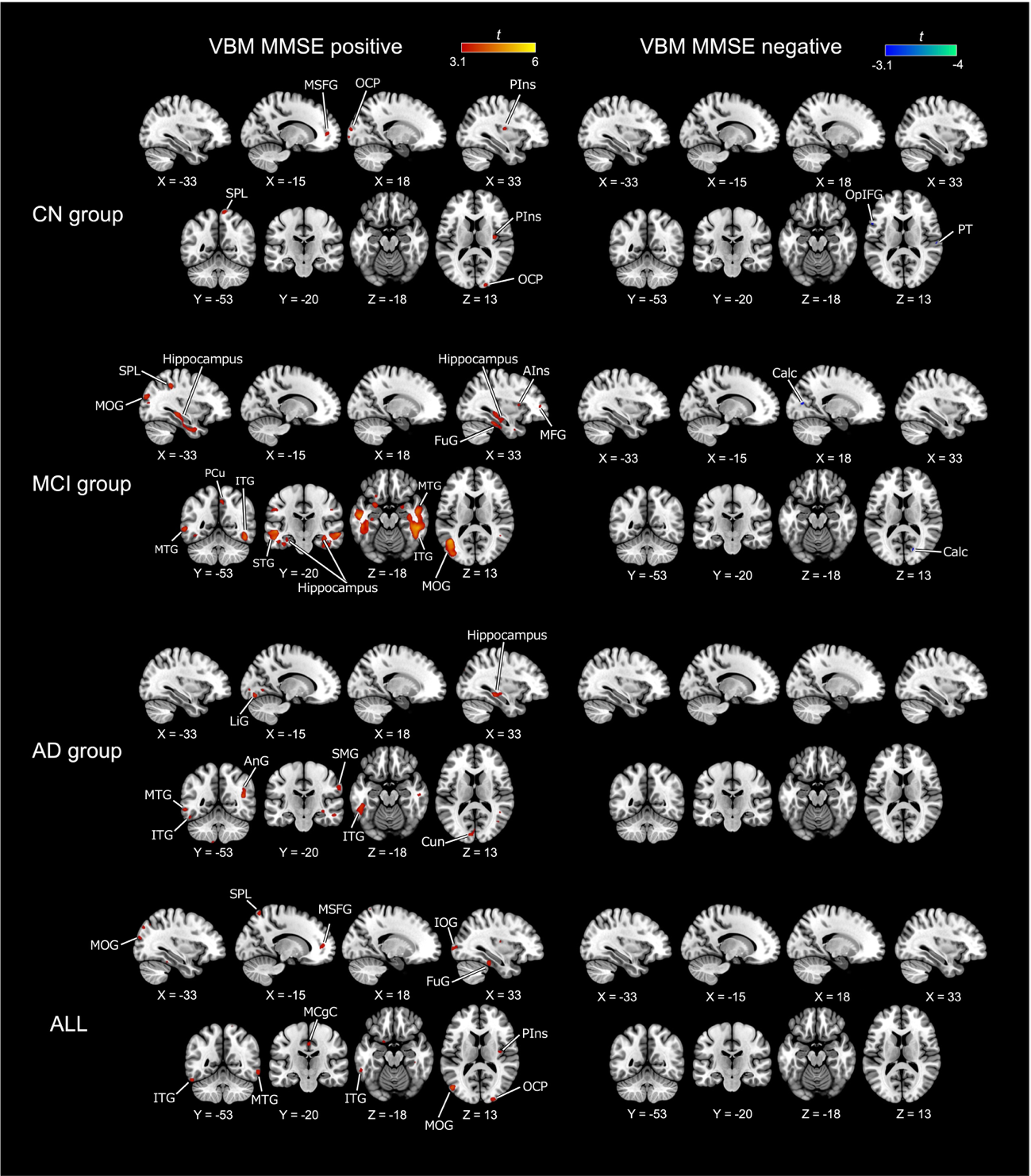

**Supplementary Figure 2. Voxel-wise associations between MMSE scores and gray matter volume across diagnostic groups.**

Regions showing positive (warm colors) and negative (cool colors) associations with MMSE are displayed for CN, MCI, AD, and all participants. Statistical maps are shown at an uncorrected threshold for visualization.
